# In your phase: Neural phase synchronisation underlies visual imagery of faces

**DOI:** 10.1101/762062

**Authors:** Andrés Canales-Johnson, Renzo C. Lanfranco, Juan Pablo Morales, David Martínez-Pernía, Joaquín Valdés, Alejandro Ezquerro-Nassar, Álvaro Rivera-Rei, Agustín Ibanez, Srivas Chennu, Tristan A. Bekinschtein, David Huepe, Valdas Noreika

## Abstract

Mental imagery is the process through which we retrieve and recombine information from our memory to elicit the subjective impression of “seeing with the mind’s eye”. In the social domain, we imagine other individuals while recalling our encounters with them or modelling alternative social interactions in future. Many studies using imaging and neurophysiological techniques have shown several similarities in brain activity between visual imagery and visual perception, and have identified frontoparietal, occipital and temporal neural components of visual imagery. However, the neural connectivity between these regions during visual imagery of socially relevant stimuli have not been studied. Here we used electroencephalography to investigate neural connectivity and its dynamics between frontal, parietal, occipital and temporal electrodes during visual imagery of faces. We found that voluntary visual imagery of faces is associated with long-range phase synchronisation in the gamma frequency range between frontoparietal electrode pairs and between occipitoparietal electrode pairs. In contrast, no effect of imagery was observed in the connectivity between occipitotemporal electrode pairs. Gamma range synchronisation between occipitoparietal electrode pairs predicted subjective ratings of the contour definition of imagined faces. Furthermore, we found that visual imagery of faces is associated with an increase of short-range frontal synchronisation in the theta frequency range, which temporally preceded the long-range increase in the gamma synchronisation. We speculate that the local frontal synchrony in the theta frequency range might be associated with an effortful top-down mnemonic reactivation of faces. In contrast, the long-range connectivity in the gamma frequency range along the fronto-parieto-occipital axis might be related to the endogenous binding and subjective clarity of facial visual features.

## 1. Introduction

Visual mental imagery—or simply visual imagery—is the process through which we retrieve and recombine information from the long-term memory, resulting in the subjective impression of “seeing with the mind’s eye”^1^. Identification of the neural processes underlying imagery is essential both for the understanding of cognition under normal circumstances^2–4^ and for the understanding and treatment of certain psychiatric disorders, such as schizophrenia and posttraumatic stress disorder^5–7^. Furthermore, several lines of research emphasise the integral role of visual imagery in social cognition^8,9^. For instance, individuals who can generate vivid visual images of other persons also form more accurate social memories^10^. On the other hand, the superior ability of visual pictorial imagery is positively correlated with social anxiety^11^. In contrast, individuals with an autism spectrum condition marked by social communication difficulties exhibit less efficient visual imagery despite having superior visuoconstructional abilities compared to neurotypical individuals^12^.

While visual imagery and visual perception share critical neural processes and regions^13,14^, their interaction and spatial overlap are not uniform. In particular, researchers have suggested that sensory networks may be engaged differently by visual imagery and perception^15,16^. While the sensory input to the primary sensory regions drives visual perception with a subsequent engagement of the secondary sensory regions and frontoparietal networks, a reverse hierarchy seems to underlie voluntary visual imagery with frontal regions executing top-down reactivation of memories and sensory components^17^. Ganis & Schendan (2008) investigated whether visual perception and visual imagery of faces recruit the same early perceptual processes, as assessed by the event-related potentials (ERPs) N170 and vertex positive potential (VPP) using a rapid adaptation paradigm^18^. As expected, face stimuli elicited a robust N170 potential localised to the posterior occipitotemporal scalp (as well as a VPP at the central site). However, while perceived adaptors (physical stimuli) diminished ERP amplitude, visualised adaptors (imagined stimuli) enhanced it, suggesting that perceived adaptors modulate perception via bottom-up mechanisms, whereas visualised adaptors affect it via top-down mechanisms. We propose that dynamic top-down interactions between cortical sites may drive two fundamental components of visual imagery - working memory and feature binding.

Top-down reactivation and active maintenance of visual images involve multiple mnemonic processes, e.g. when we imagine a known face, we need to recall and maintain it in our working memory for the duration of a mental image. It has been shown that memory retrieval and working memory are core processes underlying visual imagery^19,20^. While working memory is primarily associated with activation of regions within the frontal cortex^21^, especially in the theta frequency range^22–24^, working memory of visual mental objects is also associated with increased high-frequency activity in the gamma range (> 30 Hz) in parietal brain regions^24–29^ (also see^30^). Furthermore, the visuospatial working memory of mental images involves a broader neural network of occipital and temporal visual areas under executive frontal lobe control^19^. One of the functional roles of such long-range dynamic interactions between cortical regions could be visual feature binding of sensory images reactivated and stored in working memory. Indeed, visual feature binding involves several cortical regions, including the fusiform gyrus, the dorsal premotor cortex, and several structures within the parietal cortex, as demonstrated by multiple fMRI, MEG, and TMS studies^27,31–34^. Binding of specifically facial features recruits the bilateral middle fusiform gyri and the left precuneus^35,36^. Given a multitude of cortical regions involved in the binding of facial features, a coordinated interaction between them is arguably required. Indeed, Kottlow et al. (2012) reported a link between an increase in gamma synchronisation and feature binding during a face integration task^36^.

Taken as a whole, working memory and feature binding literature suggest that dynamic network interactions might underlie generation, retrieval, and maintenance of visual mental images, such as faces. However, studies that have addressed the brain network dynamics of visual imagery are still scarce. de Borst et al. (2012), for instance, used a combination of EEG and fMRI to investigate large-scale networks involved in the visual imagery of complex scenes and found that beta and theta synchronisation in frontal regions preceded activation of parietal and occipital regions relative to an auditory control task^37^. This finding led the authors to propose a model of mental imagery with a “frontal relay station” orchestrating visual imagery of scenes. In a more recent magnetoencephalography (MEG) study, Dijkstra et al. (2018) explored temporal dynamics of category representation (faces and houses) during visual perception and visual imagery^38^. Perception was characterised by high temporal specificity and distinct processing stages, whereas imagery showed wide generalisation and low temporal specificity from the onset of stimulus/image. Furthermore, an overlap between perception and imagery was observed around 130 ms, which decreased around 210 ms, and increased again from 300 ms after stimulus/image onset. It seems that it took a longer time to generate visual representations of mental imagery based on purely top-down processes that either did not rely on early perceptual representations, or such representations were transient and variable over time.

Here, we characterise the brain dynamics of visual imagery of faces by using EEG phase-synchrony and spectral power analysis. Based on studies reporting theta phase coherence in the frontal lobes as a key EEG signature of working memory^19,39,40^, we predicted that visual imagery of faces would be associated with frontal interhemispheric synchronisation in the theta frequency range. Furthermore, given that visual imagery is driven by top-down modulation of sensory regions^18,41^, whereas central maintenance and binding of visual representations involve gamma synchronisation^21^, we predicted that visual imagery of faces would be associated with gamma phase synchronisation along the midline fronto-parieto-occipital network. Finally, given that visual imagery involves activation of the occipital and temporal regions along the ventral stream^1,7,42^, we predicted that connectivity of the occipital lobe would extend to the occipitotemporal gamma synchronisation. Altogether, we hypothesised that the temporal dynamics of visual imagery of faces would involve three distinguishable neural processes: (i) frontal theta synchronisation, (ii) frontoparietal and occipitoparietal gamma synchronisation, (iii) and occipitotemporal gamma synchronisation.

## 2. Methods

### 2.1. Participants

Twenty-one right-handed participants (12 male; mean age = 21.20 years) with normal or corrected-to-normal vision, no history of psychiatric or neurological disorders, and no current use of any psychoactive medications, took part in the experiment. All signed an informed consent accepting their participation in the study in accordance with the Declaration of Helsinki and approved by the institutional ethics committee of the Faculty of Psychology of Universidad Diego Portales (Chile).

### 2.2. Stimuli

Stimuli were 10 greyscale images of faces, 1 greyscale image of an oval, and 10 words, each presented on a black background. All face images were full-front views of celebrities with clothing, background, and hair removed (for examples, see Fig. 1 and Supplementary Fig. 1). The word stimuli were the 10 names of their respective celebrities, presented in a black background in Helvetica font. Face images (7.32 × 8.87 cm) were presented inside an oval that subtended ∼4.19 × 5.08° of visual angle. Oval-shaped stimuli have been used previously in face perception ^43–49^ and face imagery ^50,51^studies, as they eliminate irrelevant inter-stimuli variance in low-level visual features of background. Participants sat 100 cm from the monitor. For a description of stimuli validation, see Supplementary Materials, Section A. Stimuli were presented on a 17-inch LCD monitor using Python and Pygame software.

**Figure 1.**
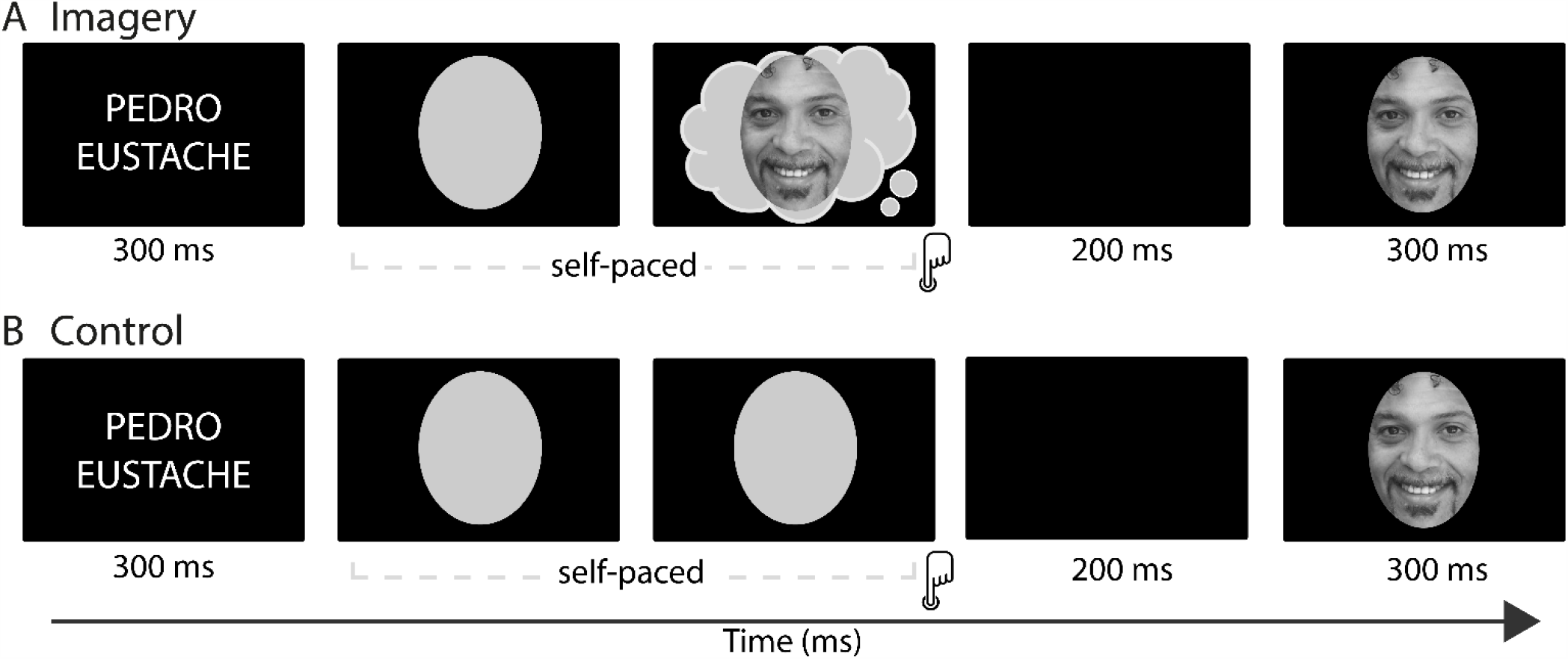
Diagram of an experimental trial for the visual imagery and control conditions. Visual imagery and control conditions had a parallel structure. **(A)** In the visual imagery condition, a celebrity face was visualised mentally after seeing the corresponding name that was presented for 300 ms. Participants pressed a key as soon as they had generated a vivid mental image, and 200 ms after this keypress, a test face appeared for 300 ms which was of the same person whose face has been mentally visualised before. **(B)** In the control condition, after seeing the name of a celebrity, participants were asked to press a key after seeing a grey oval on the screen when they felt the urge to do so. Then, 200 ms after the keypress, a test face appeared (300 ms) which was congruent with the name of the celebrity presented before.

### 2.3. Procedure

We used a modified version of the paradigm developed by Ganis & Schendan (2008)^18^ with three experimental blocks: control condition, visual imagery practice, and visual imagery condition. In the control condition (Fig. 1B), after presentation of a name of one out of ten celebrities for 300 ms, a grey oval was presented on the centre of the screen. Participants were asked to press a key after seeing a grey oval when they felt the urge to do so. Then, 200 ms after the keypress, a test face appeared (300 ms) which was congruent with the name of the celebrity presented before. There were 240 trials in the control condition, with 24 trials per each celebrity name. The imagery practice started with participants memorising pictures of 10 faces of a new set of celebrities with 12 exposures for each. Next, participants practised mentally visualising each memorised image three times. During the imagery practice block, the name appeared followed by a black screen with a grey oval centred in the middle of the screen where participants visualised the picture of the named face. Having done so, they pressed a key to see the actual picture to adjust their mental image. In the imagery condition (Fig. 1A), the name of a celebrity appeared for 300 ms, followed by a black screen with a middle-centred grey oval where subjects mentally visualised the associated memorised picture. Participants were instructed to generate a mental image of the face cued by the celebrity’s name shown and to project that mental image onto the grey oval on the screen. They were instructed to press a key once the mental image was very clear. They were given a maximum of 10 seconds for a keypress. In a case of no response, the experiment moved to the next trial. 200 milliseconds after the keypress the test picture appeared for 300 ms. For all trials, the test picture was the picture of the face that was visualised mentally. There were 240 trials in the imagery condition, with 24 trials per each celebrity name. After the imagery condition, participants were asked to rate the vividness, contour definition, and emotional expression of the images on a 1-10 Likert scale.

### 2.4. Electrophysiological recordings and pre-processing

EEG signals were recorded with 129-channel saline-based HydroCel sensor nets using a GES300 Electrical Geodesic amplifier at a rate of 500 Hz, and NetStation software. The physical filters were set at 0.01–100 Hz for the recording acquisition, and a notch filter at 50 Hz was applied offline for removing the DC component. During acquisition, Cz electrode was used as a reference. For the analyses, the electrodes’ average was used as the reference electrode. Two bipolar derivations were designed to monitor vertical and horizontal ocular movements. Eye movement contamination and other artefacts were removed from data before further processing using independent component analysis^52^. Continuous data were epoched from −1500 to 0 ms around the button press (0 ms).

Trials that contained voltage fluctuations exceeding ± 200 μV, transients exceeding ± 100 μV, or electrooculogram activity exceeding ± 70 μV were rejected. After screening, no significant differences in the number of trials were observed between conditions. Each participant yielded at least 91% of artefact-free trials which represented a minimum of 210 trials per condition. To address the sample size bias imposed by our phase-locking measure (see below), we used the same number of trials for all conditions and participants by randomly selecting 210 artefact-free trials in those conditions that exceeded this minimum. The EEGLAB MATLAB toolbox was used for data pre-processing and pruning^52^.

### 2.5. Analysis of endogenous EEG activity related to visual imagery

There is growing evidence that endogenous or ‘‘ongoing’’ neural activity, i.e. activity that is not time-locked to stimuli, is functionally significant^30^. A classical experimental approach for studying endogenous changes in the EEG related to internal neural fluctuations during cognitive tasks is to analyse the EEG window *before* the onset of motor responses when participants report internal changes. This approach has previously been used for studying neural signatures of bistable awareness^53–55^, binocular rivalry^56^ and intrusions of consciousness^57^. Following the same rationale, but this time for the analysis of visually *imagined* faces, we computed ongoing EEG activity preceding the onset of each response (button press), i.e. response-locked activity. We performed two separate analyses: one that captures local synchrony (spectral power) and the second analysis that captures long-range synchrony (phase synchrony) between signals in relation to visual imagery (described below in full).

To determine the length of the EEG window of analysis, we identified the trial with the shortest reaction time relative to the onset of the grey oval for both the control (1760 ms) and the visual imagery conditions (2870 ms). Our criteria for making conditions comparable was that the grey oval should be present during the entire window of analysis. Thus, we decided to use a window of 1500 ms for all trials and conditions (from −1500 to 0 ms locked to the response time). Time-frequency charts of spectral power and phase-synchrony are expressed in z-scores relative to the beginning of this period (−1500 to −1250 ms), which was regarded as a baseline.

### 2.6. Between-electrodes measure: long-range phase synchronisation

We quantified phase locking between pairs of electrodes to measure dynamical interactions among electrode signals oscillating in the same frequency range. Phase synchronisation analysis proceeds in two steps: (i) the estimation of the instantaneous phases and (ii) the quantification of the phase locking.

#### 2.6.1. Estimation of the instantaneous phases

To obtain the instantaneous phases, *φ*, of the neural signals, we used the Hilbert transform approach^58^. The analytic signal *ξ*(*t*) of the univariate measure *x*(*t*) is a complex function of continuous time defined as

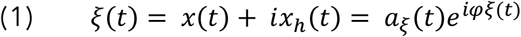

where the function is the Hilbert transform of

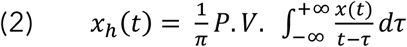

P.V. indicates that the integral is taken in the sense of Cauchy principal value. Sequences of digitised values give a trajectory of the tip of a vector rotating counter-clockwise in the complex plane with elapsed time.

The vector norm at each digitising step *t* is the state variable for instantaneous amplitude *a*_*ξ*_(*t*). This amplitude corresponds to the length of the vector specified by the real and imaginary part of the complex vector computed by Pythagoras’ law and is equivalent to the magnitude of the observed oscillation at a given time and frequency point.

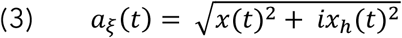

is the arctangent of the angle of the vector with respect to the real axis and is the state variable for instantaneous phase *φ*_*x*_(*t*).

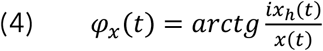

The instantaneous phase *φ*_*x*_(*t*) of *x*(*t*) is taken to be equal to *φ*_*ξ*_(*t*). Identically, the phase *φ*_*y*_(*t*) is estimated from *y*(*t*). This phase is thus the angle of the vector specified by the real and imaginary components. For a given time and frequency point, it corresponds to a position inside the oscillation cycle (peak, valley, rising, or falling slope).

The instantaneous phase, although defined uniquely for any kind of signal to which the Hilbert transform can be applied, is challenging to interpret physiologically for broadband signals. For this reason, a standard procedure is to consider only narrow-band phase synchronisation by estimating an instantaneous phase for successive frequency bands, which are defined by band-pass filtering the time series^59^. Thus, we band-pass filtered EEG signals in multiple consecutive 1 Hz-wide frequency bins from 1 to 60 Hz using a zero-phase shift non-causal finite impulse filter.

#### 2.6.2. Phase locking quantification: weighted phase lag index (wPLI)

Phase synchronisation can be considered as an EEG measure of information exchange between neuronal populations^60^. Generally speaking, phase synchronisation is calculated from the phase or the imaginary component of the complex cross-spectrum between signals measured at a pair of channels. For example, the well-known Phase Locking Value (PLV; see^61^) is obtained by averaging the exponential magnitude of the imaginary component of the cross-spectrum. However, many of such phase coherence indices derived from EEG data are affected by the volume conduction^62^. As a result, a single dipolar source can produce spurious coherence between spatially disparate EEG channels, rather than the values being related to a pair of distinct interacting sources per se. The Phase Lag Index (PLI), first proposed by Stam et al. (2007), attempts to minimise the impact of volume conduction and common sources inherent in EEG data by averaging the signs of phase differences, thereby ignoring average phase differences of 0 or 180 degrees^63^. This is based on the rationale that such phase differences are likely to be generated by volume conduction of single dipolar sources. Despite being insensitive to volume conduction, PLI has two important limitations: first, there is a strong discontinuity in the measure, which causes it to be maximally sensitive to noise; second, when calculated on small samples, PLI is biased toward strong coherences (i.e. it has a positive sample-size bias). Formally, the PLI is defined as the absolute value of the sum of the signs of the imaginary part of the complex cross-spectral density *Sxy* of two real-valued signals *x*(*t*) and *y*(*t*) at time point or trial *t*:

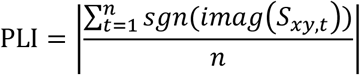

The Weighted PLI measure (wPLI; see^64^) addresses the former problem by weighting the signs of the imaginary components by their absolute magnitudes. The PLI sample size problem can be addressed by considering the same amount of trials per condition in the phase coherence analysis. Further, as the calculation of wPLI also normalises the weighted sum of signs of the imaginary components by the average of their absolute magnitudes, it represents a dimensionless measure of connectivity that is not directly influenced by differences in spectral or cross-spectral power. Thus, wPLI addresses potential confounds caused by volume conduction, by scaling contributions of angle differences according to their distance from the real axis, as almost ‘almost-zero-lag’ interactions are considered as noise affecting real zero-lag interactions:

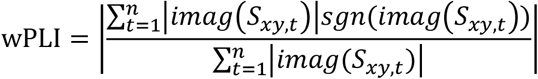

For these reasons, we employed wPLI to estimate connectivity in our data (see Supplementary Fig. 2). The wPLI index ranges from 0 to 1, with a value of 1 indicating perfect synchronisation (phase difference is perfectly constant throughout the trials) and value 0 representing the total absence of synchrony (phase differences are random). wPLI charts are expressed in z-scores (standard deviation units) relative to the beginning of this period (−1500 to −1250 ms), which was regarded as a baseline.

### 2.7. Within-electrodes measure: spectral power

Given that phase synchronisation metrics depend on the amplitude of EEG oscillations^64^, a hypothetical increase of local spectral power in frequencies of interest may confound cross-electrode synchronisation changes. To control for it, spectral power was estimated by calculating the square of the envelope obtained from the absolute value of the Hilbert transform after filtering. This procedure provides instantaneous power emission, which reflects the strength of local synchronisation. The frequency bands were defined as follows: theta (4–7 Hz) and gamma (30–60 Hz), following the canonical frequency-band classification (e.g. ^65^).

### 2.8. Selection of electrodes of interest

Groups of electrodes were selected for spectral power analysis (within-electrodes measure) and phase synchrony analysis (between-electrodes measure) by selecting canonical frontal, parietal, occipital and temporal electrodes. In the case of the spectral gamma power analysis, power values within frontal and parietal electrodes were averaged per condition and participant. Similarly, for spectral theta power analysis, values within frontal electrodes were averaged per condition and participant. In the case of long-range frontoparietal, occipitoparietal and occipitotemporal wPLI analysis, we calculated the mean connectivity that every electrode of one group shared with every electrode of another group. In such a case, connectivity values within each electrode group were discarded from the analysis. In the case of interhemispheric fronto-frontal wPLI analysis, we calculated the mean connectivity that every electrode of the right-frontal hemisphere shared with every electrode of the left-frontal hemisphere discarding connectivity values between right-frontal electrodes and between left-frontal electrodes. This procedure allowed to test specifically the role of long-distance interactions during visual imagery. Finally, wPLI values of the corresponding regions of interest were averaged per condition.

### 2.9. Statistical analysis

For EEG results, a cluster-based nonparametric statistical framework implemented in FieldTrip^66^ was used throughout the analysis of the power and wPLI time-frequency charts. In brief, time-frequency windows of interest were compared in pairs of experimental conditions (control vs. imagery). For each such pairwise comparison, epochs in each condition were averaged subject-wise. These averages were passed to the analysis procedure of FieldTrip, the full details of which are described elsewhere^67^. In short, this procedure compared corresponding time-frequency points in the subject-wise averages using one-tailed dependent (for within-subject comparisons) or independent (for between-subject comparisons) samples *t*-tests. Although this step was parametric, FieldTrip uses a nonparametric clustering method to address the multiple comparisons problem. *t* values of adjacent time-frequency points whose *P* values were lower than 0.05 were clustered together by summating their *t* values, and the largest such cluster was retained. This whole procedure, i.e. calculation of *t* values at each time-frequency point followed by clustering of adjacent *t* values, was then repeated 1000 times, with recombination and randomised resampling of the subject-wise averages before each repetition. This Monte Carlo method generated a nonparametric estimate of the *p*-value representing the statistical significance of the initially identified cluster. The cluster-level *t* value was calculated as the sum of the individual *t* values at the points within the cluster.

Multiple and simple regression analyses were performed using MATLAB (2016a; MathWorks, Inc.) and Jamovi (Version 0.8.1.6) [Computer Software] (Retrieved from https://www.jamovi.org; open source) statistical software.

## 3. Results

### 3.1. Behavioural analysis and scales

Regarding reaction times, pairwise comparisons revealed a significant difference in reaction times between visual imagery (M = 4603, SD = 1043 ms) and control (M = 2550 ms, SD = 614 ms) conditions (t(1,20) = 8.83; p<0.001). After imagery condition, participants were asked to rate the vividness (M = 2.76, SD = 0.944), contour definition (M = 3.05, SD = 0.865), and emotional expression (M = 2.10, SD = 0.889) of the images on a 1-5 Likert scale.

### 3.2. Gamma phase synchrony patterns related to imagery

On frontoparietal electrode pairs (Fig. 2D), phase synchronisation in the gamma range (30-60 Hz) was significantly higher in the visual imagery condition compared to the control condition (Fig. 2). Cluster-based nonparametric analysis showed a significant cluster from −346 to −194 ms in the gamma range (34-60 Hz; *p* = 0.018; Fig. 2C). Interestingly, no significant clusters were found between conditions in the spectral power of the gamma frequency range in this time-frequency window (Supplementary Fig. 3).

**Figure 2.**
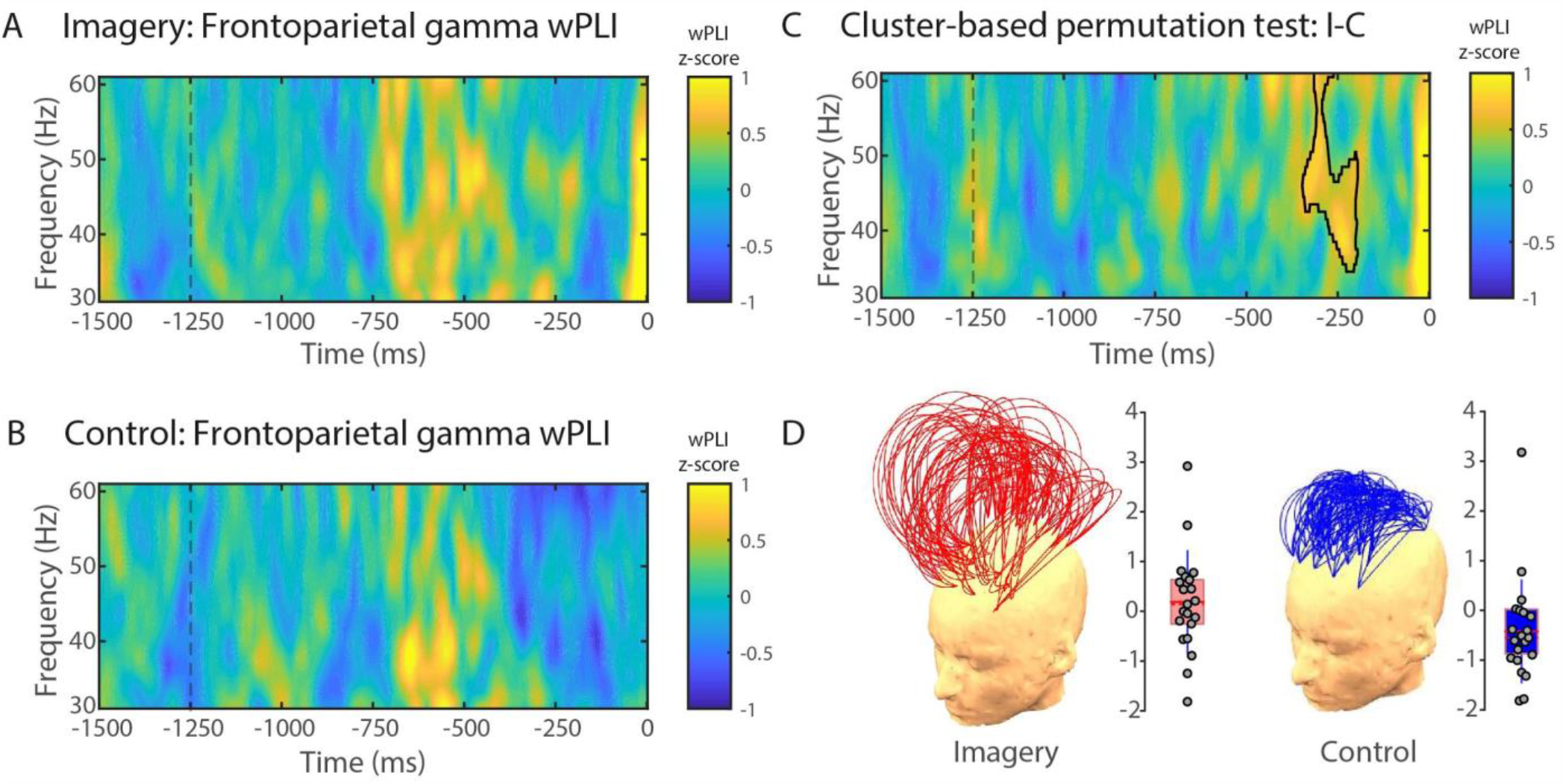
Gamma phase-synchrony between frontal and parietal electrodes. wPLI (30-60 Hz) for wPLI cluster between conditions (visual imagery minus control) in the gamma band (30-60 Hz). **(D)** Region of interest (ROI) for phase-synchrony analysis. wPLI values are expressed in standard deviations (z-scores) in reference to the baseline (−1500 to −1250 ms). Trial length (−1500 ms) is relative to response time (0 ms). **(D)** Topographical representation of frontoparietal wPLI channel pairs and single-participant wPLI values for the imagery (left) and control (right) conditions. Each arc represents a pair of channels, and the height of the arc is its normalised value. Grey circles represent single-participant wPLI values for the cluster depicted in **(C)** for the imagery (left panel) and control (right panel) conditions. The red horizontal line represents the group mean, the rectangle represents SEM, and the red-dashed horizontal line represents the group median.

Similarly to the frontoparietal connectivity, phase synchronisation in the gamma frequency range (30-60 Hz) was significantly higher between the occipitoparietal electrode pairs in the visual imagery condition compared to the control condition (Fig. 3). The cluster-based nonparametric analysis showed a significant cluster from −268 to −186 ms in the gamma range (36-60 Hz; *p* = 0.012; Fig. 3C). Contrary to this, there was no significant difference in the occipitotemporal gamma phase synchronisation between the visual imagery and control conditions.

**Figure 3.**
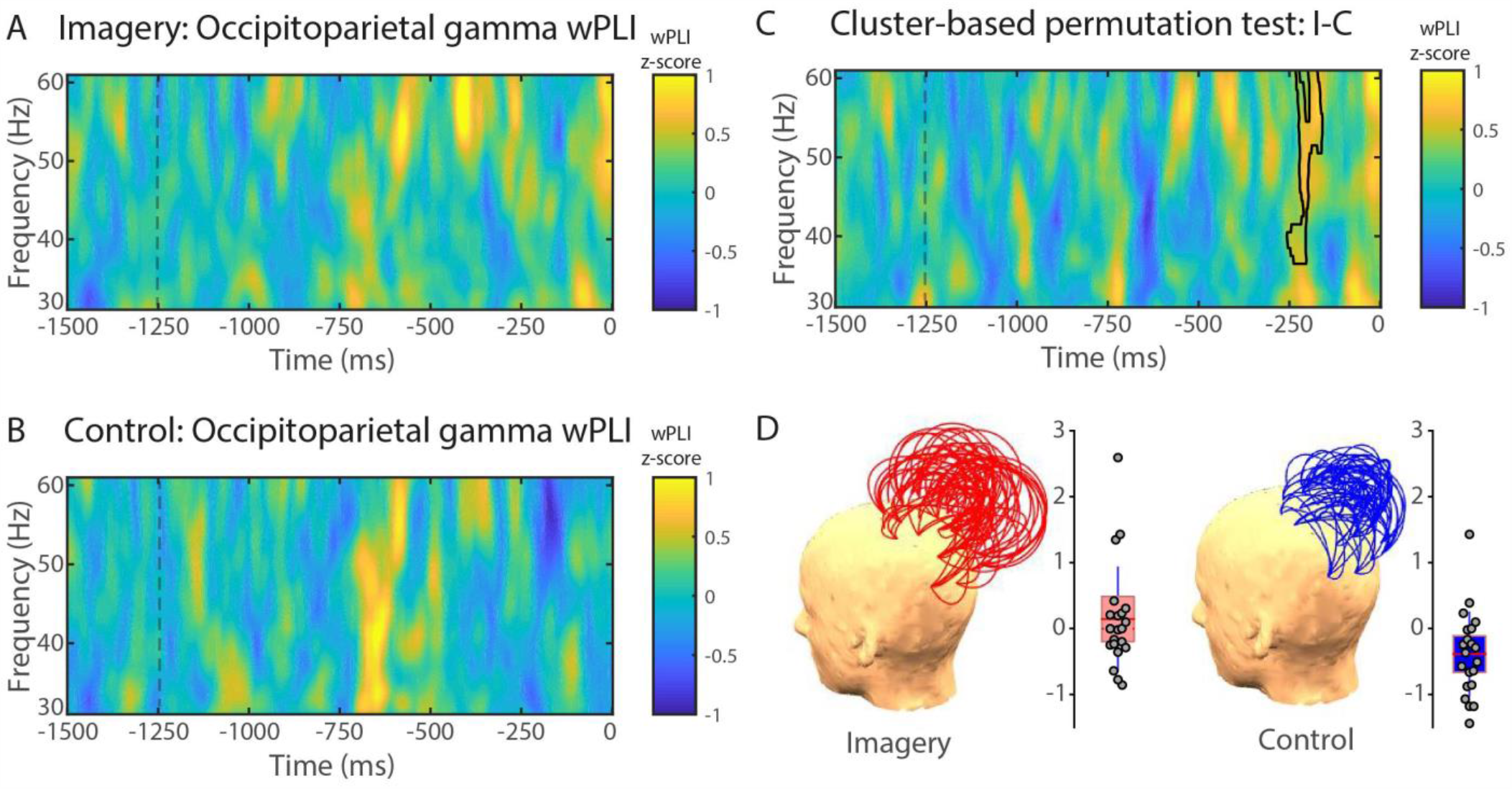
Gamma phase-synchrony between occipital and parietal electrodes. wPLI (30-60 Hz) for wPLI cluster between conditions (visual imagery minus control) in the gamma band (30-60 Hz). **(D)** Region of interest (ROI) for phase-synchrony analysis. wPLI values are expressed in standard deviations (z-scores) in reference to the baseline (−1500 to −1250 ms). Trial length (−1500 ms) is relative to response time (0 ms). **(D)** Topographical representation of occipitoparietal wPLI channel pairs and single-participant wPLI values for the imagery (left) and control (right) conditions. Each arc represents a pair of channels, and the height of the arc is its normalised value. Grey circles represent single-participant wPLI values for the cluster depicted in **(C)** for the imagery (left panel) and control (right panel) conditions. The red horizontal line represents the group mean, the rectangle represents SEM, and the red-dashed horizontal line represents the group median.

### 3.3. Theta phase synchrony patterns related to imagery

Phase synchronisation in the theta range (5-7 Hz) increased in the imagery condition compared to the control condition from −998 to −330 ms between interhemispheric frontofrontal pairs of electrodes (*p* = 0.019; Fig. 4). No significant clusters were found between conditions in the spectral power of theta frequency range in this time-frequency window (Supplementary Fig. 4).

**Figure 4.**
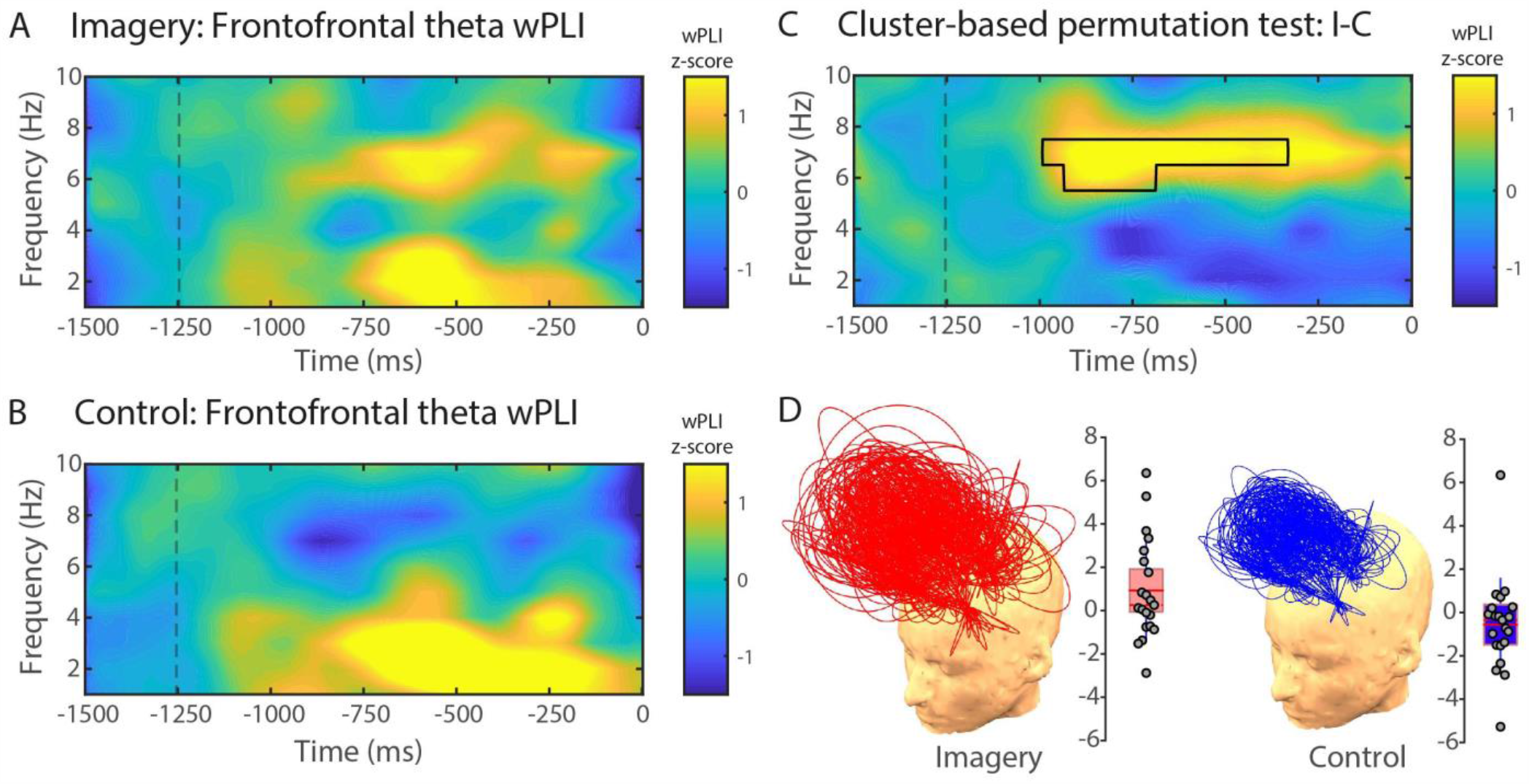
Theta phase-synchrony between inter-hemispheric frontal electrode pairs. wPLI (1-10 Hz) for visual imagery **(A)** and control **(B)** conditions. **(C)** Cluster-based permutation test comparing visual imagery and control conditions. The area highlighted in the time-frequency chart depicts a significant wPLI cluster between conditions (visual imagery minus control) in the theta band (5-7 Hz). **(D)** Region of interest (ROI) for phase-synchrony analysis. Only interhemispheric frontofrontal pairs of channels were considered for the analysis (see Section 2.8 for details). wPLI values are expressed in standard deviations (z-scores) in reference to the baseline (−1500 to −1250 ms). Trial length (−1500 ms) is relative to response time (0 ms). **(D)** Topographical representation of frontofrontal wPLI channel pairs and single-participant wPLI values for the imagery (left) and control (right) conditions. Each arc represents a pair of channels, and the height of the arc is its normalised value. Grey circles represent single-participant wPLI values for the cluster depicted in **(C)** for the imagery (left panel) and control (right panel) conditions. The red horizontal line represents the group mean, the rectangle represents SEM, and the red-dashed horizontal line represents the group median.

### 3.4. Relationship between gamma phase-synchrony and behavioural scales

We further investigated, in an exploratory manner, the statistical dependencies between phase synchronisation in the gamma band, and the subjective report of imagined faces in three different dimensions. We performed separate multiple regressions using vividness, contour definition and emotional valence Likert scales as outcome variables, and frontoparietal and occipitoparietal gamma-band wPLI as predictors (Figure 5).

**Figure 5.**
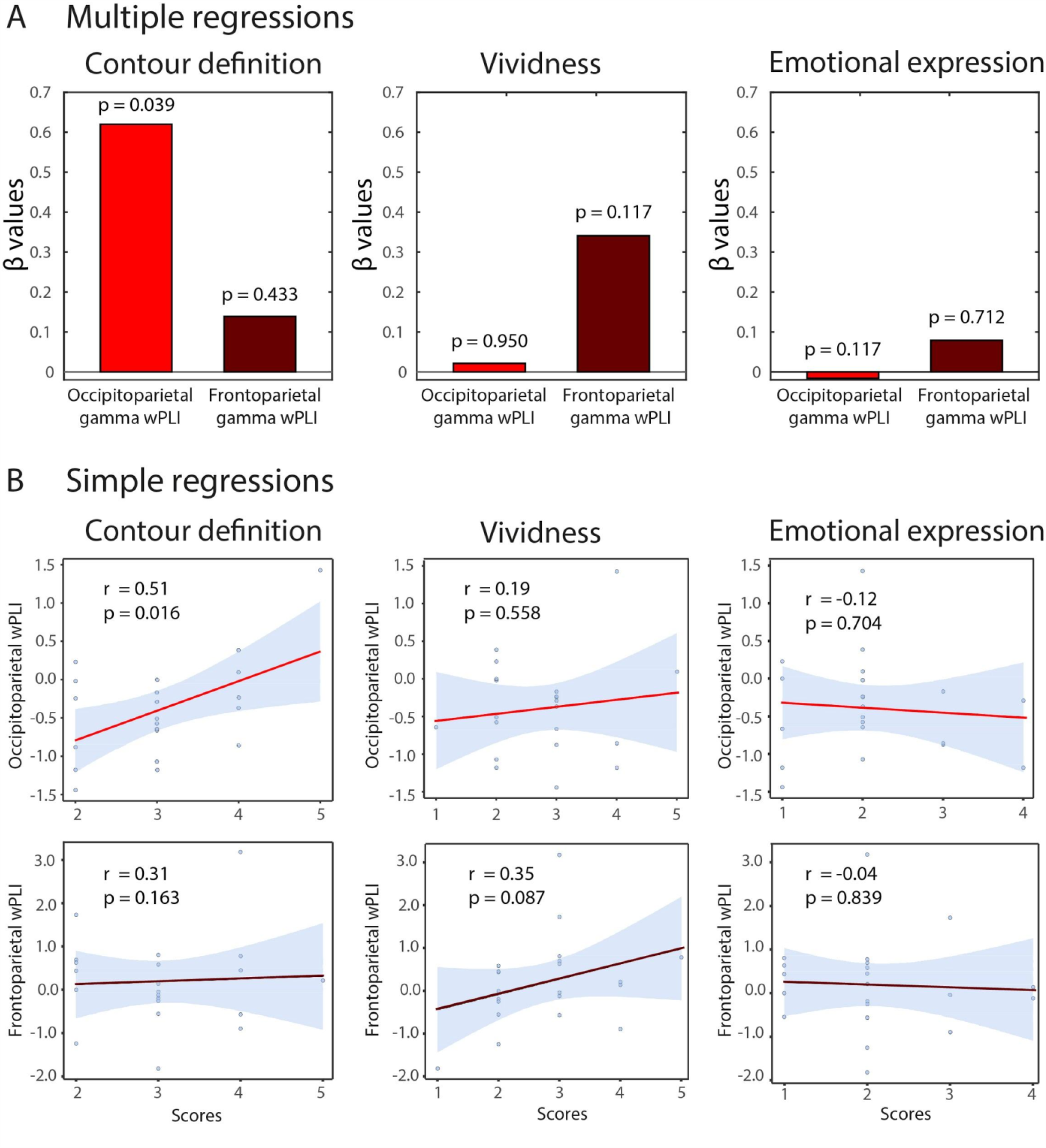
Multiple and single regressions between wPLI and behavioural scales. **(A)** Beta coefficients for three separate multiple regressions using Likert scales (vividness, contour definition, and emotional expression) as an outcome variable and frontoparietal and occipitoparietal wPLI as predictors. **(B)** Pearson’s correlation between frontoparietal and occipitoparietal wPLI values and the corresponding scale scores.

In the Contour Definition model (R^2^ = 0.29; F(2,18) = 3.74; p=0.044), occipitoparietal gamma-band wPLI predicted the contour definition scores of imagined faces (β=0.620.; p=0.039), while frontoparietal gamma-band wPLI was not a reliable predictor (β=0.139; p=0.433). However, in the Vividness model (R^2^ = 0.14; F(2,18) = 1.55; p=0.240), none of the wPLI indices predicted vividness of the imagined faces (frontoparietal gamma-band wPLI: β=0.034; p=0.117; occipitoparietal gamma-band wPLI: β=0.02; p=0.950). Similarly, in the Emotional Expression model (R^2^ = 0.14; F(2,18) = 1.55; p=0.240), none of the wPLI indices predicted emotional valence of the imagined faces (frontoparietal gamma-band wPLI: β=-0.07; p=0.576; occipitoparietal gamma-band wPLI: β=-0.02; p=0.950). This relationship was also described in terms of a simple Pearson’s correlation, showing a significant association between contour definition scores and occipitoparietal gamma-band wPLI (r=0.51; p=0.016) but not for gamma-band frontoparietal wPLI (r=0.31; p=0.163) (Fig 5B).

## 5. Discussion

Successful retrieval from long term memory and effective feature integration, are essential processes enabling us to visualise a familiar face. Here we show that visual imagery of faces involves long-range EEG phase synchronisation. When contrasted with a control condition of seeing an oval figure, the face imagery task ignited (1) theta phase inter-hemispheric synchronisation between frontal electrodes within a time window from −998 to −330 ms and (2) gamma phase synchronisation along the fronto-parieto-occipital axis within a time window from −346 to −186 ms before participants reported their mental image was subjectively clear. Arguably, these neural signatures could be linked to the mnemonic reactivation of visual features of known faces and subsequent binding of reactivated facial features, respectively.

In particular, the increase in the synchronisation of the theta frequency range between frontal electrodes during the imagery task may be related to top-down reactivation of face memories^68,69^. Given that the fronto-parieto-occipital gamma synchronisation temporally followed the offset of the frontal theta effect, we suggest that gamma synchronisation may be related to the endogenous binding of facial visual features transiently sustained in working memory. Such binding may be essential for the clarity of a sustained image and eventual recognition of the imagined face. Indeed, we found that occipitoparietal synchronisation was associated with the subjective perception of contours of imagined faces, suggesting its role in the generation of coherent mental images with subjectively clear boundaries. This finding complements previous reports that gamma activity in the occipitoparietal region is associated with visual feature binding of Kanizsa-type illusory contours^70^. However, our cognitive interpretation of neural findings is based on previous studies about the mechanisms of visual representations in the working memory^20,21,30,71^ and binding of facial features^36,72^. Given that mental imagery and control conditions did not include specific manipulations of visual working memory load or feature binding, we recognise the speculative nature of this interpretation.

Notably, it has been shown that induced gamma activity in scalp EEG can reflect micro-saccadic eye movements^73^, while visual imagery tasks typically facilitate involuntary saccades^74^. To control for the possibility that increased gamma phase synchronisation in the imagery condition might have been due to eye movements, spectral power analysis was carried out for gamma frequencies in the same time window. The absence of gamma power differences between conditions suggests that long-range synchronisation results cannot be attributed to eye movements.

It is worth noting that a number of studies have reported that changes in gamma or theta power and its temporal dynamics are associated with visual binding or episodic memory tasks^68,69,71,72,75,76^. In particular, there have been reports of increased power (i.e. local synchronisation) in the beta and gamma frequencies during visual binding. Melloni et al. (2007), for example, found that this increase occurs specifically during preconscious processing stages of visual input in a bottom-up manner and therefore does not necessarily result in conscious detection^75^. We hypothesise that because mental imagery is a top-down, consciously driven process^16,41^, it is not accompanied by the changes in local power that characterise other visual tasks. A complementary hypothesis is that the apparent discrepancy in gamma power findings between perceptual tasks and imagery tasks may reflect the neural distinction between exogenously driven and endogenously driven binding processes. Such a hypothesis would predict that the neural signature of endogenous binding is not gamma range intensity (i.e. power), but cross-channel synchronisation.

There is increasing evidence that mental imagery involves top-down activation of both dorsal and ventral pathways of visual information processing^17^. In such case, when imagining faces of known people, the ventral stream, especially fusiform face area, would be involved in the reactivation of facial features, whereas the dorsal stream would bind them into a spatially coherent whole (for a review, see^77^). We thus hypothesised that the imagery of faces would be associated with long-range gamma synchronisation between both the occipitoparietal electrode pairs covering the dorsal stream and the occipitotemporal electrode pairs covering the ventral stream. However, we found no evidence that the occipitotemporal gamma synchronisation is associated with visual imagery of faces. Nevertheless, given that EEG spatial specificity is very low^62,78^, we do not exclude the possibility that our methodology was not sensitive enough to detect ventral stream activation. It is also possible that the temporal electrodes selected for EEG analysis were too anterior to the fusiform face area, or perhaps that signal-to-noise ratio in the electrodes chosen was too low to detect high-frequency phase locking.

This study was concerned with the neural mechanisms underlying visual imagery of faces, including those that overlap with normal face perception. Thus, our control condition was devised so that it did not include a face as the test stimulus but instead featured a grey oval, identical to the one employed in the imagery condition. This meant that both conditions consisted of the same stimuli with the manipulation being only in the instructions given to participants in each condition. It could be argued that the results can be explained by a repetition effect and/or a memory effect rather than by imagery, as the imagery practice session always preceded the imagery condition. However, when stimuli are repeated, neural activity is usually reduced^79^. For instance, gamma frequency range power (> 30 Hz) decreases following repetition of familiar objects across lags of one or two intervening objects^26^. On the contrary, active retention and encoding of one or more items in working memory have generally been associated with an increase in theta and gamma power^80–82^. Thus, theta and gamma amplitudes could be expected to decrease due to the repetition effect, whereas an increase in the amplitude of these frequency bands in the imagery condition relative to visual perception could be expected due to the memory effect. However, both possibilities were excluded because theta and gamma amplitudes did not change during the time windows where phase-synchrony increased. Alternatively, the repetition and memory effects on spectral power cancelled each other. Future studies on the neural dynamics of visual imagery should address this matter directly.

Our study presents several limitations. First, our results indicate dynamic changes with topographic specificity; however, we could not reliably infer brain regions generating these effects. Future studies should source-reconstruct the EEG signals to link neural dynamic findings to brain sources. Second, our interpretation requires reverse inference based on past studies and temporal occurrence, given the straightforward approach of our visual imagery task. Future studies should include additional behavioural manipulations that can allow distinguishing between memory retrieval, memory maintenance, and binding of facial features. Finally, we cannot discard the possibility that some participants may have experienced spontaneous mental images during the control condition. In such a case, the reported EEG findings could arguably reflect cognitive control and effort associated with task instructions for the imagery condition. However, the results suggest that the influence of spontaneous imagery, if any, was not decisive. First, response times in the control condition were much shorter than in the imagery condition. Mental imagery is an effortful process that requires longer processing time^38,83,84^. Second, we found several predicted EEG synchronisation differences between the control and imagery conditions. The lower theta and gamma phase synchronisation during the control condition suggests that whatever spontaneous mental images were produced, they should have been much less complex than images generated by the effortful visual recall. Third, we found that occipitoparietal gamma synchronisation predicted the strength of subjectively reported contour definition of imagined faces, linking the observed neural dynamics with subjective quality of visual images. Nevertheless, while our results demonstrate differences in neurodynamics between the control and imagery conditions, future studies should come up with additional instructions or a secondary task that would prevent spontaneous imagery from occurring. On this point, it would be relevant to collect subjective reports of spontaneous imagery after control blocks.

Overall, the present study indicates that the imagery of faces is associated with spatially broad phase synchronisation in the theta and gamma frequency ranges, arguably involving face memory-retrieval and visual feature binding.

## AUTHOR CONTRIBUTIONS STATEMENT

A.C-J. conceived the study; A.C.J., R.C.L., A.R-R., S.C. and V.N. developed the methodology; A.C.J., R.C.L., D.M-P., J.P.M., J.V., collected the experimental data; A.C-J., R.C.L., A.E.-N., T.A.B., S.C., interpreted the findings; A.C-J., R.C.L., V.N., A.I., wrote the manuscript; A.E-N., D.H., constructively reviewed the manuscript.

