## Supplemental Material for "In your phase: Neural phase synchronisation underlies visual imagery of faces"

### SUPPLEMENTARY MATERIALS

#### A. Stimuli Validation

In the following, we describe the validation and standardization of stimuli suitable for visual imagery research in a Chilean population using Ganis and Schendan's (2008) paradigm.

#### PARTICIPANTS

A total of 60 undergraduate students, aged 18-36 years (M= 26.11; SD= 3.21), participated in the validation study – all from the cities of Santiago, Viña del Mar and Valparaíso, Chile. They were asked to participate voluntarily either directly by the researchers or by public ads posted on their universities' bulletin boards.

#### STIMULI VALIDATION

**Faces validation and recognition.** To validate facial images, we administered a questionnaire including the images of 125 celebrities (55 females, 70 males). Participants were asked to say the name of each celebrity. Faces were considered valid when responses reached a minimum of 90% correct identification by the group tested. 55% of the pictures (i.e. 70) met this criterion and were thus selected to be used in the main study.

As the pictures were selected randomly, the recognition of the celebrities might have been affected by participants' age. In order to test that, we performed an ANOVA with age as a categorical factor that includes the following groups: ages 18-24 years (M= 16.12, SD= 5.16); ages 25-30 years (M= 16.53, SD= 3.83); ages 31-36 years (M= 16.75, SD= 3.33). No significant differences were found between these groups [$F(2.372) = 0.71$; $p = 0.48$], suggesting that participants' age did not affect celebrity recognition.

### STIMULUS NORMALIZATION

**Faces.** Seventy images of validated celebrity faces were subsequently normalized based on size, brightness, and light intensity. Next, they were converted to a grayscale image using Adobe® Photoshop®. In order to avoid potential distractions, only regions corresponding to eyes, nose and mouth were selected, keeping these facial areas inside an oval, as symmetrically as it was possible for each image (Supplementary Fig. 1).

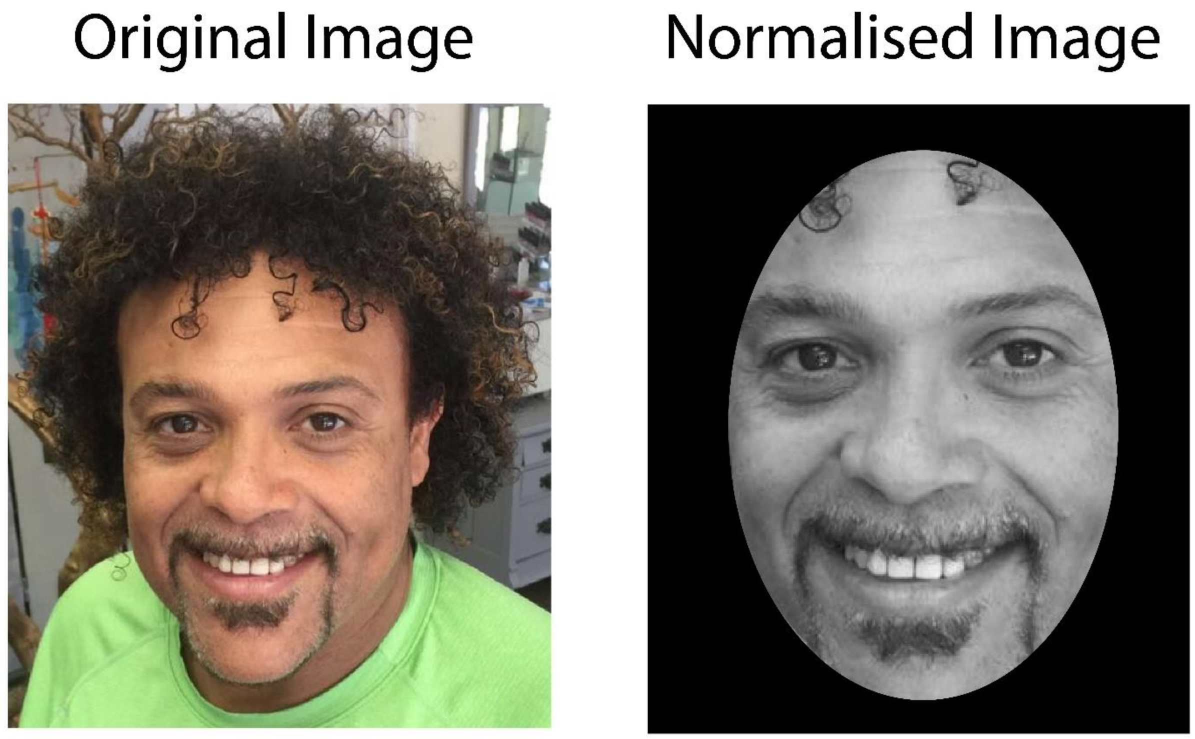


**Supplementary Figure 1.** Face photo normalization. Typical celebrity photo used in the recognition test: the left image shows an example of those presented during the validation phase; the image on the right shows a normalized face ready to be used in the experimental task.

### STANDARDIZED STIMULUS VALIDATION

**Standardized face normalization.** In order to determine whether the modifications described above affected image recognition, another recognition test was applied to a different set of 25 participants (M= 23.11, SD= 1.51; 14 female). As before, faces that were recognized by 90% of participants were considered as valid. All images met this criterion, and hence no image was rejected. In addition, an interclass ANOVA was applied to examine the extent to which image variability affected face recognition variability. The result (0.05) indicated that only 5% of face recognition variability could be attributed to image variability after the normalization process.

**Emotional valence recognition.** We tested whether the emotional valence of each face image was positive. Each face was classified within one of the following categories of emotional valence: positive (M= 93.97, SD= 7.59), neutral (M= %.85, SD= 7.49) or negative (M= 0.34, SD= 1.64). An ANOVA showed a significant main effect of category ($F(2.204)= 4901.48$; $p= 0.00032$)). Post-hoc analysis (Tukey HSD test, MS = 190.330,58, DF= 2.0) showed that the means of the faces with positive valence (M=93.62, SD= 1.06) were greater than faces with either neutral valence (M= 88.11, SD= 1.06; $p<0.01$) or with negative valence (M= 93.62, SD= 1.06; P<0.01). Recognition was considered valid as above 94% of the images were classified as having a positive emotional valence. Finally, we tested whether celebrity gender (i.e. male or female) affected the recognition of faces with positive valence. An ANOVA did not show a significant effect of gender ($F(1,1723)= 0.41$; $p=0.52$), suggesting that gender does not affect facial recognition of faces with positive valence.

### B. Supplementary Figures

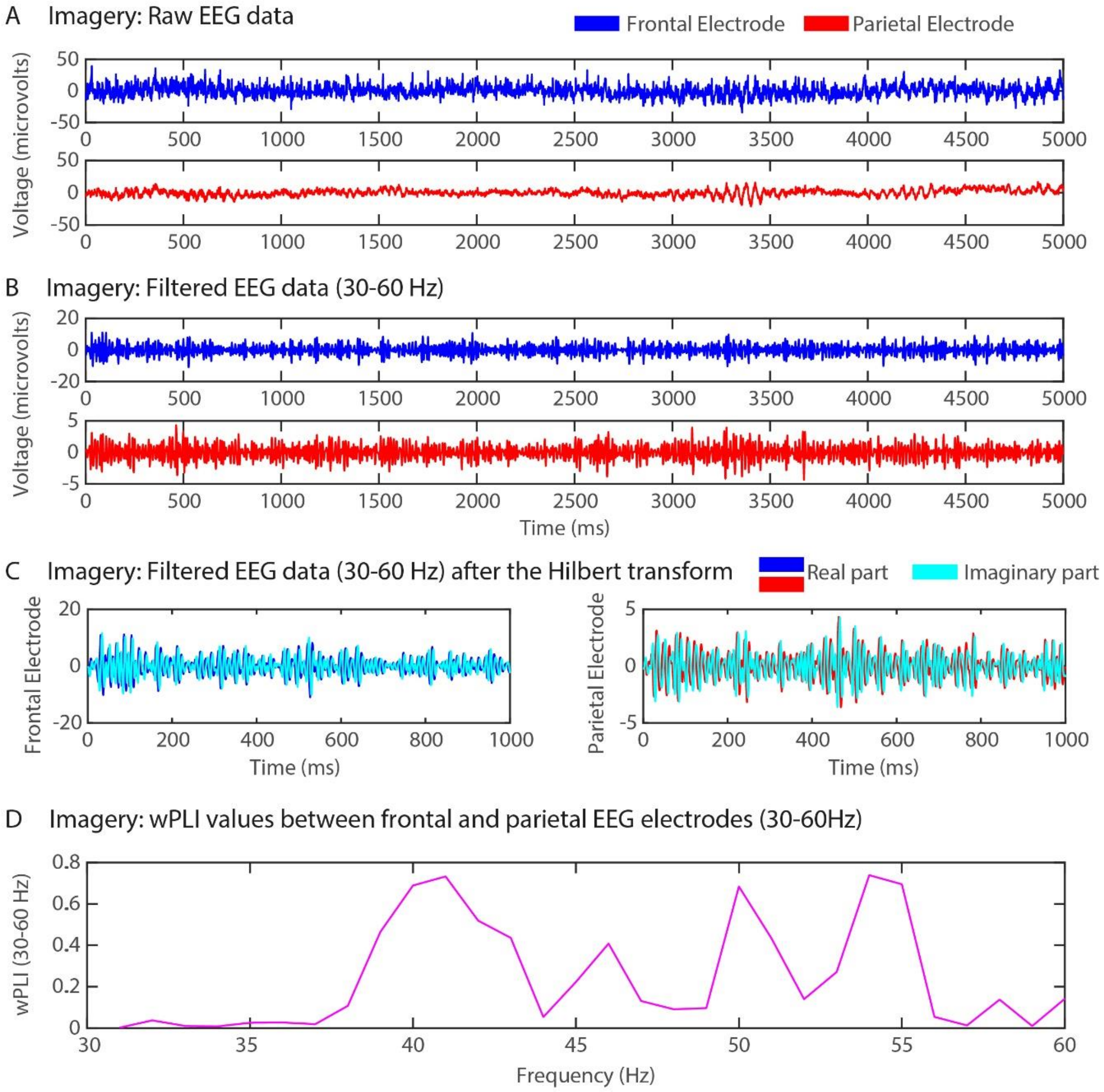


**Supplementary Figure 2.** Schematic EEG processing pipeline. **(A)** 5-second raw EEG trace for a frontal (blue) and a parietal electrode (red) in the imagery condition of a selected participant. **(B)** Same as (A) but after band-pass filtering in the gamma frequency band (30-60 Hz) and after a notch filtering for 50 Hz line noise (50 Hz). **(C)** 1-second EEG trace (second 2 to 3 in plots A and B) after computing the Hilbert transform. The real part of the frontal and parietal electrodes is depicted in blue and red respectively, and their corresponding imaginary part is depicted in cyan. **(D)** wPLI between frontal and parietal electrodes between 30 and 60 Hz, calculated over the same 1-second period and averaged over time. wPLI values shown here are not z-scored.

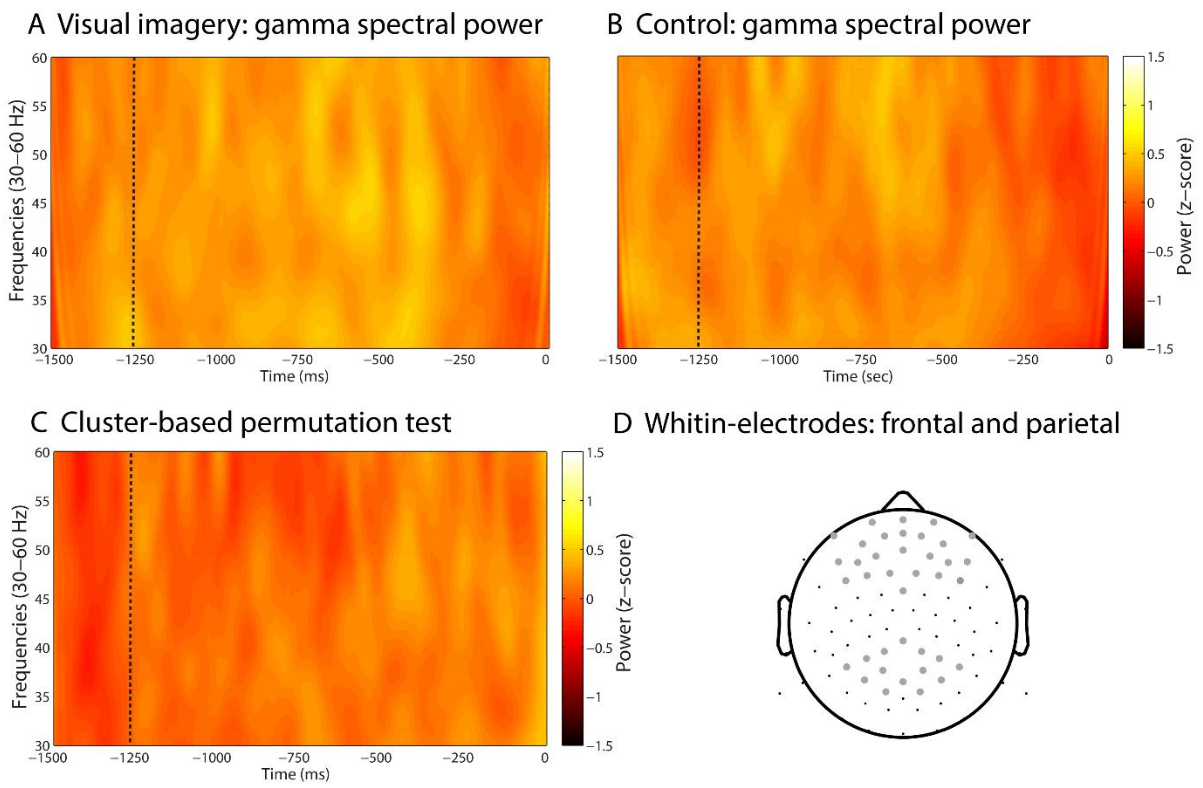


**Supplementary Figure 3.** Gamma spectral power in the frontal and parietal electrode sites**.** Gamma spectral power (30-60 Hz) for visual imagery **(A)** and control **(B)** conditions. **(C)** Cluster-based permutation test comparing visual imagery and control showed no significant cluster between conditions. All values are expressed in standard deviations (SD) in reference to the baseline (-1500 to -1250 ms). Trial length (-1500 ms) is relative to response time (0 ms).

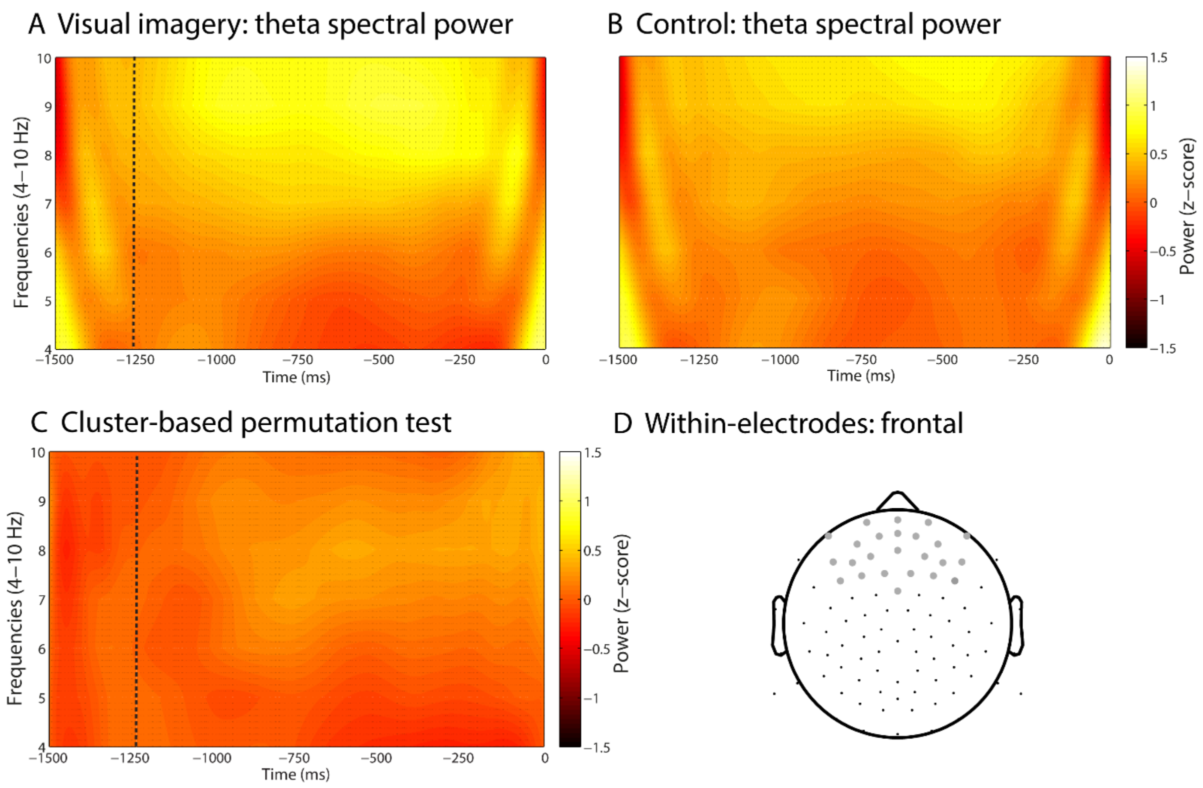


**Supplementary Figure 4.** Theta spectral power in the frontal electrode site. Spectral power (4-10 Hz) for visual imagery **(A)** and control **(B)** conditions. **(C)** Cluster-based permutation test comparing visual imagery and control revealed no significant cluster in the theta band (4-7 Hz) between conditions (visual imagery minus control). **(D)** Region of interest (ROI) for spectral power analysis. All values are expressed in standard deviations (SD) in reference to the baseline (-1500 to -1250 ms). Trial length (-1500 ms) is relative to response time (0 ms).
